# PCR artifact in testing for homologous recombination in genomic editing in zebrafish

**DOI:** 10.1101/077883

**Authors:** Minho Won, Igor B. Dawid

## Abstract

We report a PCR-induced artifact in testing for homologous recombination in zebrafish. We attempted to replace the *lnx2a* gene with a donor cassette, mediated by a TALEN induced double stranded cut. The donor construct was flanked with homology arms of about 1 kb at the 5’ and 3’ ends. Injected embryos (G0) were raised and outcrossed to wild type fish. A fraction of the progeny appeared to have undergone the desired homologous recombination, as tested by PCR using primer pairs extending from genomic DNA outside the homology region to a site within the donor cassette. However, Southern blots revealed that no recombination had taken place. We conclude that recombination happened during PCR *in vitro* between the donor integrated elsewhere in the genome and the *lnx2a* locus, as suggested by earlier work [1]. We conclude that PCR alone may be insufficient to verify homologous recombination in genome editing experiments in zebrafish.

## Introduction

Recent advances in gene and genome editing [2-6] have greatly increased the value of model systems such as the zebrafish [7-10]. Whereas introduction of deletions/insertions into a defined region of the genome of zebrafish is now routine, precise genome editing is still challenging. Multiple approaches towards this aim have been introduced, mostly by exploiting doublestranded break facilitated homologous recombination between the genome and an introduced donor DNA [11-17]. A recent publication has presented a set of procedures to achieve precise genome editing with high efficiency and accuracy [18].

In our studies of *lnx2a* gene function in pancreas development in zebrafish [19] we attempted replacement of essentially the entire gene by a donor cassette. When assayed by PCR, it appeared that our attempts had been successful. However further study, in particular using Southern blots, showed that the recombinant molecules had been generated *in vitro* and did not reflect the structure of the genome. We had not in fact achieved replacement of the gene by the donor cassette. The purpose of this paper is to summarize our experiments to issue a warning that testing for homologous recombination-mediated genome editing by PCR can be misleading, under certain circumstances. We also want to shine a spotlight on earlier observations of *in vitro* recombination during PCR reactions [1].

## Results

In earlier studies we found that the *lnx2a* gene is required for the differentiation of exocrine cells in the pancreas of zebrafish embryos [19]. During these studies we felt that it would be desirable to delete almost the entire *lnx2a* gene, a segment of about 29 kb, and replace it with a donor cassette that could assist in future studies of the locus. We designed a donor construct as illustrated in Figures 1 and 2. The construct contains a Gal4-ecdysone receptor module that would allow regulation of transcription of UAS driven transgenes, controlled by addition of the ecdysone agonist tebufemazide [20]. Further, cerulean fluorescent protein (CFP) was included (see Fig. S1)[21]. The donor cassette was flanked by homology arms of about 1 kb on the 5’ and 3’ sides, as suggested by Zu et al., 2013 [11]. We used the previously described TALEN pair, found to be efficient in generating deletions in the *lnx2a* gene [19], to create a double stranded break in the locus to stimulate recombination. In addition we introduced reagents designed to inhibit nonhomologous end joining and to stimulate homologous recombination, as suggested by Qi et al. and Panier and Boulton [22,23].

**Fig. 1.**
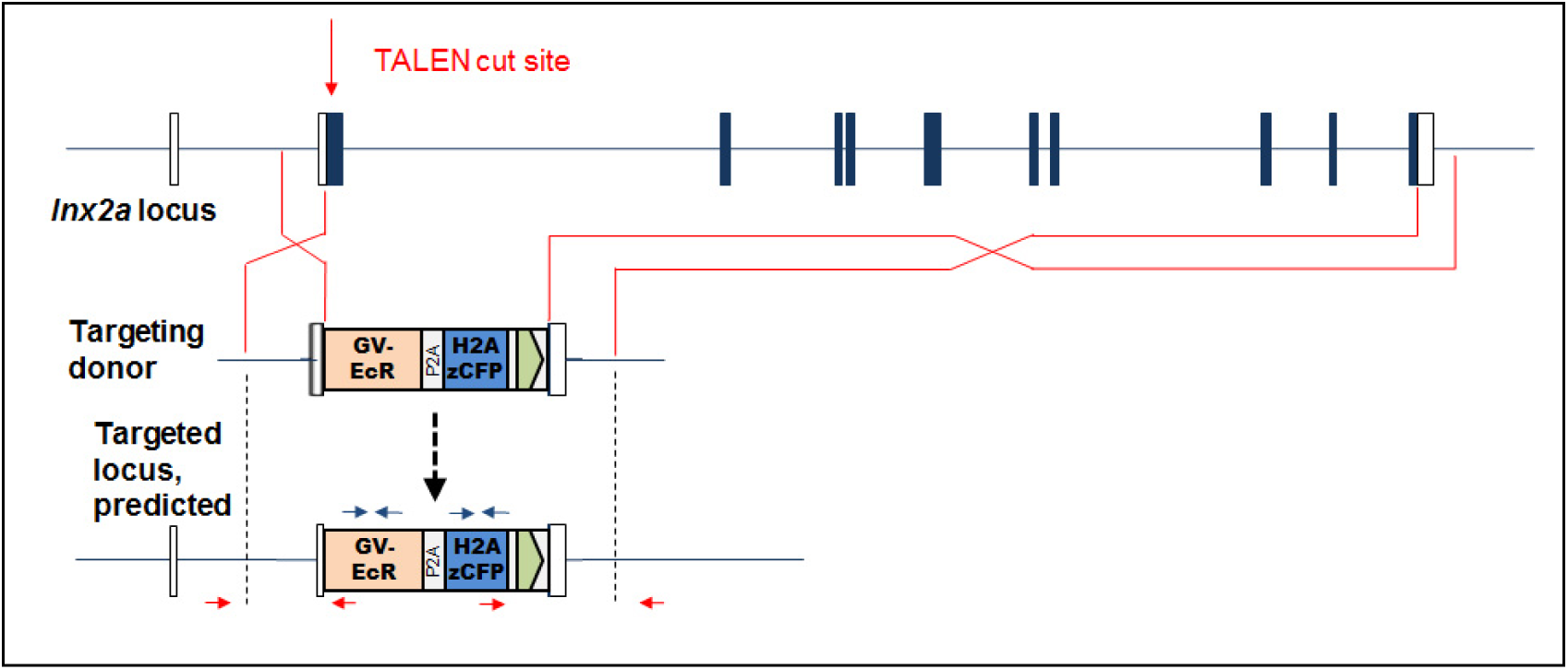
Model of ln×2a genomic locus, donor construct (abbreviated), and predicted recombination product. Small arrows indicate primer pairs. The genomic locus and donor DNA/targeted locus are shown at different scale, in this and all following figures.

**Fig. 2.**
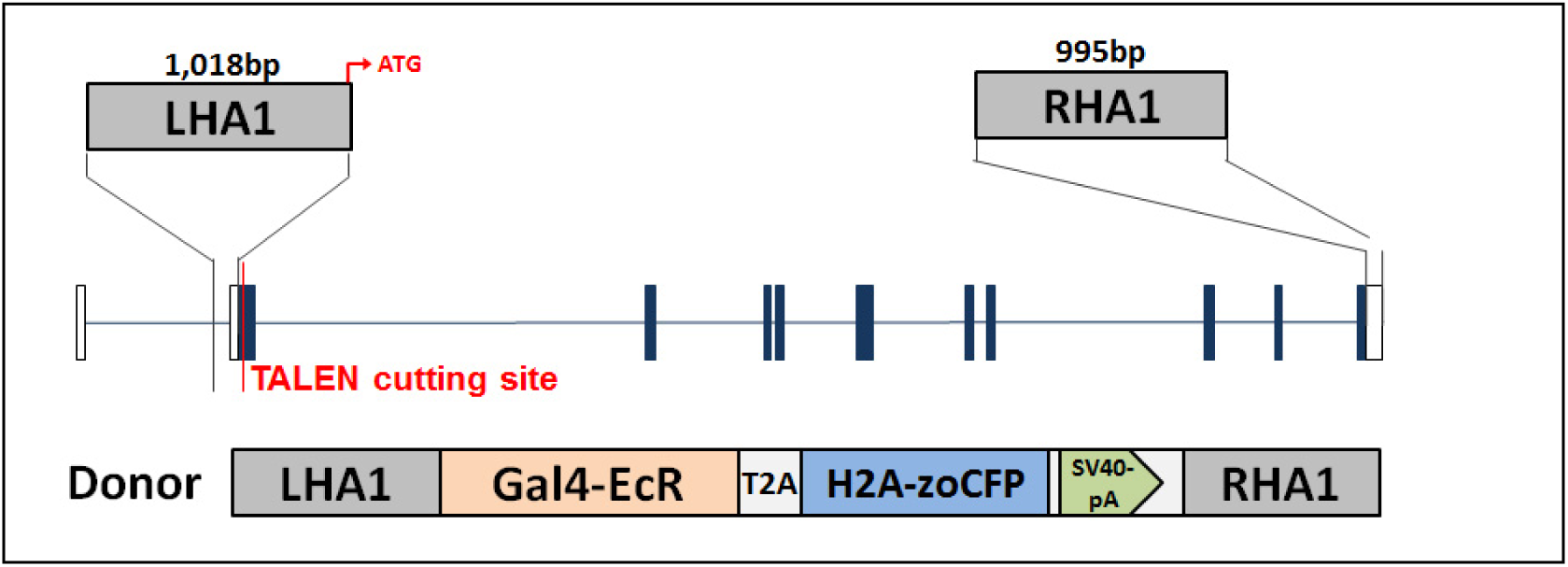
Model of Left and Right Homology arms, and complete model of donor vector. LHA1, left homology arm; RHA1, right homology arm; EcR, ecdysone receptor; zoCFP, zebrafish optimized CFP.

Zebrafish embryos were injected with reagents in various combinations. To test for recombination we used two PCR primer pairs. The 5’ pair consists of a forward primer complementary to genomic DNA upstream of the left homology arm (LHA1 in Fig. 2), and a reverse primer located within the ecdysone receptor region of the donor cassette. The 3’ pair consists of the forward primer within the CFP sequence of the donor and the reverse primer in genomic DNA downstream of the right homology arm (RHA1 in Fig. 2). The initial test used pooled DNA from 30 injected (G0) embryos in each reaction to test for occurrence of recombination in somatic cells of some of these fish. The expected PCR products were obtained with both primer pairs, with abundances that suggested increased efficiency of homologous recombination when using reagents that favored this process (Fig. 3). Encouraged by this result, we raised embryos injected under apparent optimal condition (red outline in Fig. 3B) and outcrossed them to wild type fish. Among the progeny we found multiple individuals whose DNA acted as template to produce the expected PCR products with the 5’ and 3’ primer pairs as well as with primers internal to the CFP sequence of the donor cassette (Fig. 3C).

**Fig. 3.**
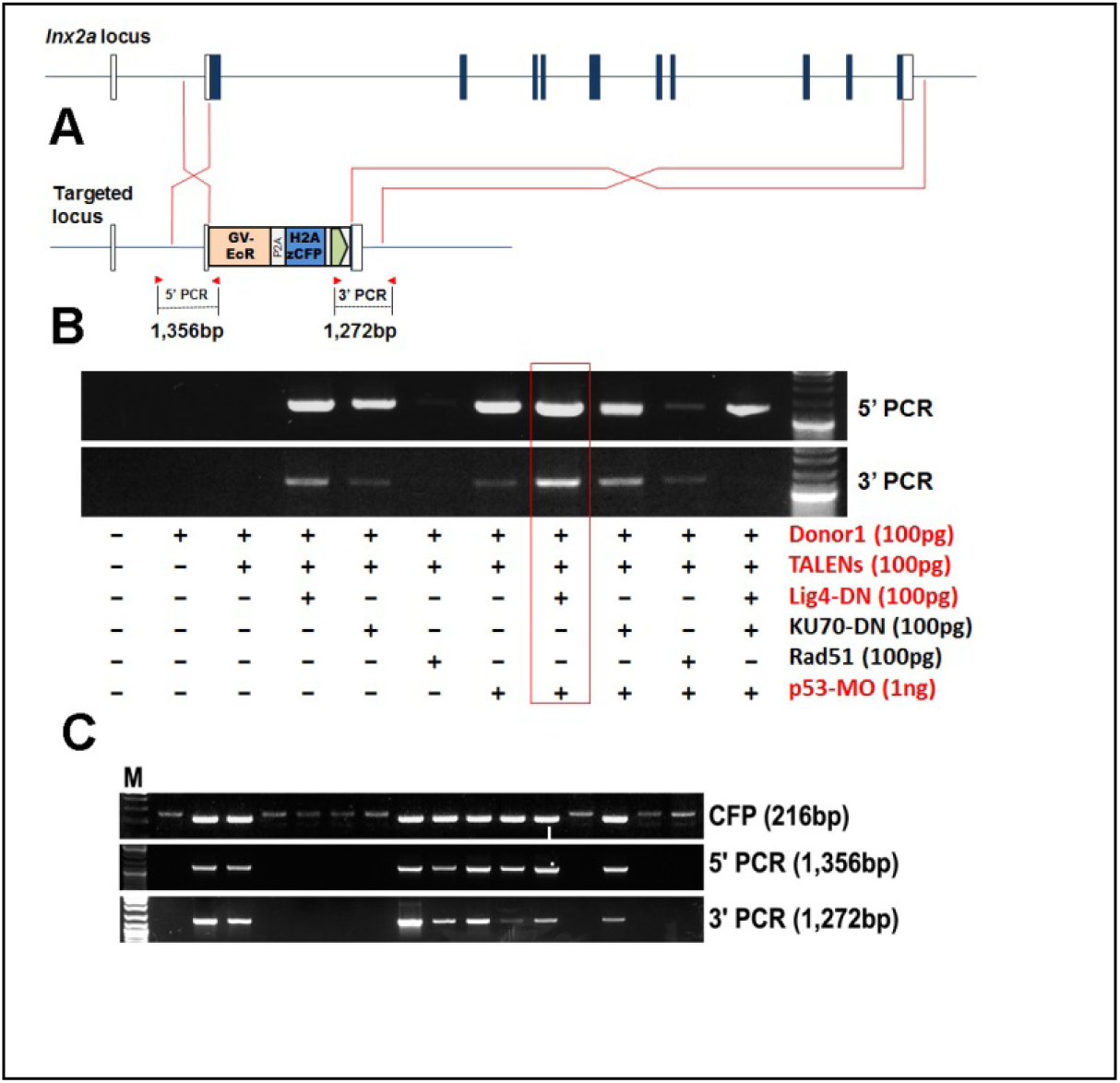
Testing for incorporation of donor DNA into the genome. (A) Model of *lnx2a* locus and predicted product of homologous recombination. (B) Groups of 30 embryos injected with the reagents specified were pooled, DNA was extracted, and PCR was performed with 5’ and 3’ primer pairs. (C) Embryos injected under optimal conditions (red outline in B) were raised and outcrossed to wild type fish. Individual progeny embryos were tested by PCR with a primer pair internal to the donor DNA (CFP) and with the 5’ and 3’ primer pairs. Shown is a selection of positive and negative embryos.

Sequencing of examples of the 5’ and 3’ PCR products revealed an excellent match to the sequence predicted for products of homologous recombination (Fig. 4). There were a number of single nucleotide differences between the sequence of the PCR product and the genome sequence, shown as red letters in Figure 4. We interpret these differences as SNPs between the sequence of the overlap region in the donor construct and the genomic sequence of the particular strain of zebrafish we used in our experiments. Under this assumption we could identify the region where the recombination had taken place; this region is outlined in red.

**Fig. 4.**
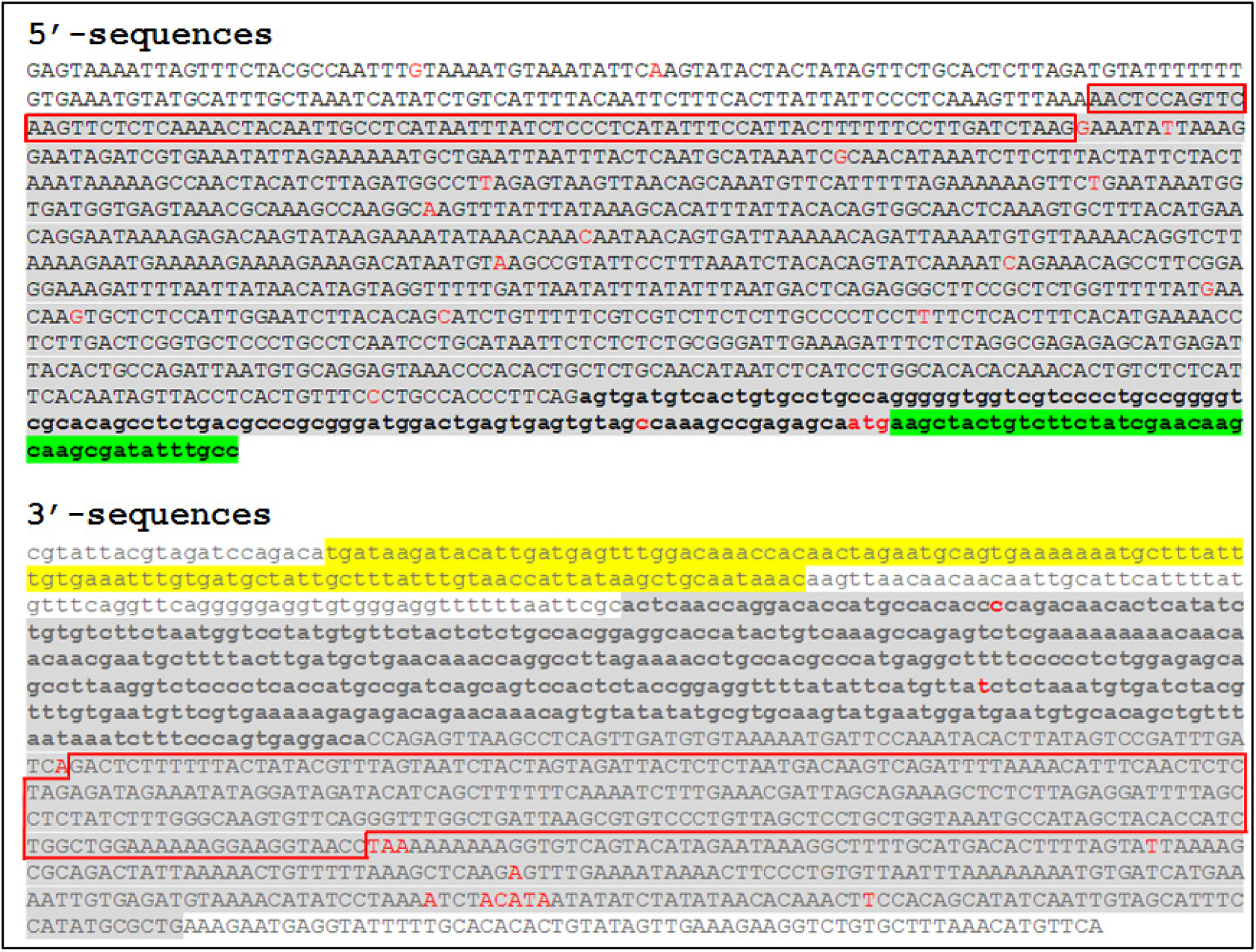
Sequence of recombinant molecules. PCR products obtained with 5’ and 3’ primer pairs (see Fig. 3C) were sequenced. Sequence without highlight is genomic sequence outside the homology (overlap) region. Grey highlight indicates genomic sequence in the homology arm. We can deduce that recombination took place in the region outlined in red because of SNPs (in red) that differ between the fish line used and the genomic clone used for donor construction. Lower case bolded sequence in the 5’ region indicates exon sequence, with red letters indicating the start codon, followed by the ecdysone receptor coding region highlighted in green. On the 3’ side, grey highlight indicates genomic sequence in the homology arm. Sequence without highlight in the 3’ end is genomic sequence outside the homology arm. Lower case letters again show exon sequence, yellow underline indicating the SV40 PA terminator sequence in the insert DNA. Nucleotides without highlight at the end of the sequence are again outside the homology arm.

While the results in Figures 3 and 4 supported the conclusion that replacement of the *lnx2a* gene by donor DNA had taken place, further studies questioned this interpretation. One indication was the fact that incrosses between putative recombinant fish should have led to 25% homozygous individuals in which the entire *lnx2a* locus was replaced. However, all progeny of such incrosses still contained the locus, as shown by PCR for internal regions. Southern blot analysis [24] provided clear evidence that locus replacement had not taken place. Figure 5 shows the structure of the predicted recombined locus, indicating the relevant Hind III sites and the probes used. Probes 1 and 2, located in the genomic region to the left of the left homology arm stained several bands and thus were not fully conclusive. However, the results seen with two additional probes were clear: neither probe revealed a 7.5 kb band as would be predicted from the recombinant locus. Probe 3, internal to the donor construct, stained a very large band in the progeny of injected fish, and nothing in the wild type. Probe 4, overlapping for just 19 nucleotides with the right homology arm and otherwise representing downstream genomic sequence, stained a 5.3 KB band with predicted size for genomic DNA in both the “recombinant” and wild type fish, as well as in a BAC clone containing the *lnx2a* locus (Fig. 5). Clearly, progeny from injected fish were transgenic for donor DNA, but the donor DNA had not replaced the *lnx2a* locus, and this locus was undisturbed.

**Fig. 5.**
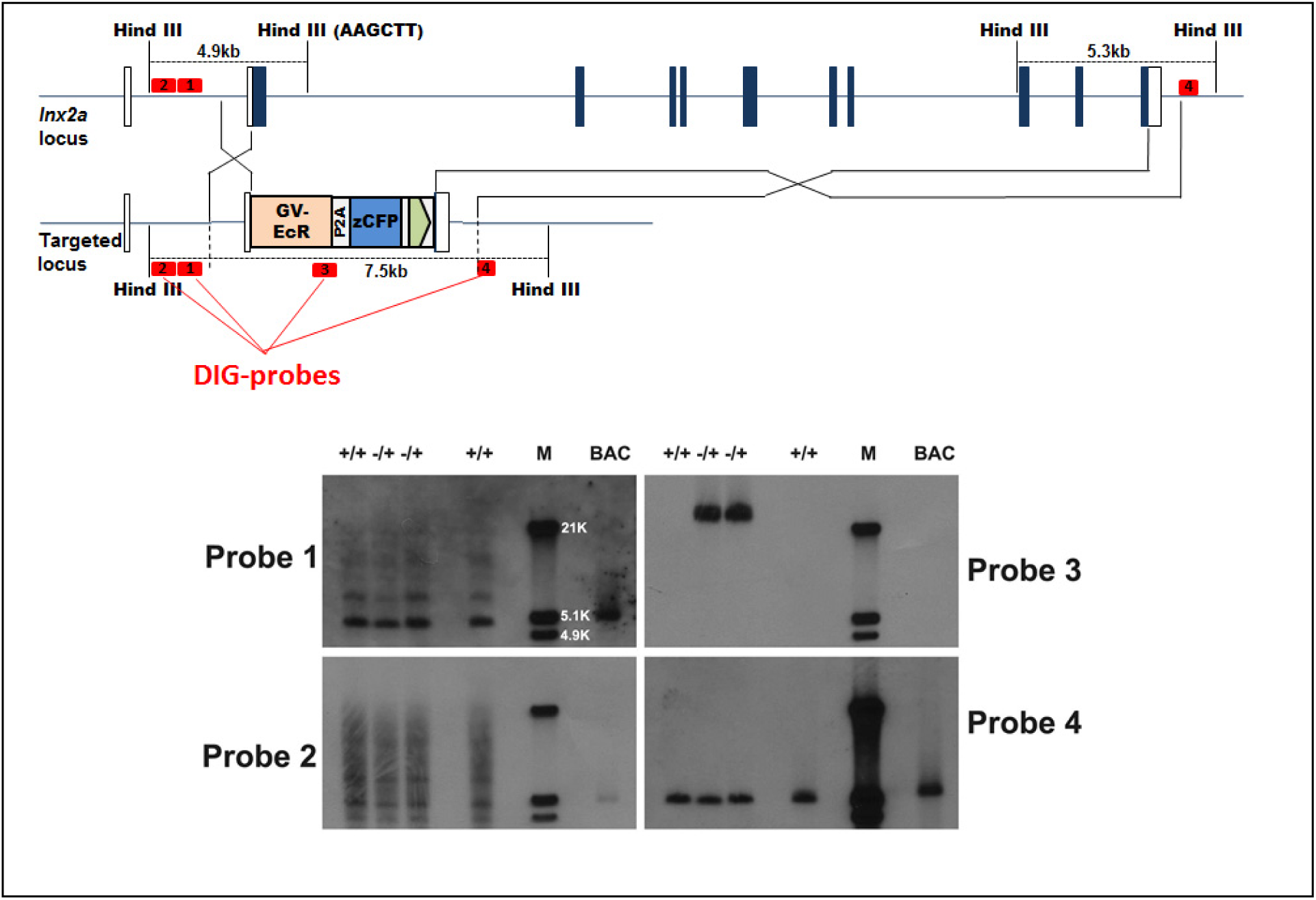
Southern blots contradict the interpretation that donor DNA was incorporated into the genome by homologous recombination. The predicted targeted locus is shown with the location of Hind III sites. The predicted recombined locus should yield a 7.5 KB Hind III fragment detectable by each of the four probes used. While bands close to 7.5 KB were seen with probes 1 and 2 in addition to other bands, probes 3 and 4 did not detect any band close to that size. The individual fish in lanes labeled +/- were predicted to be heterozygous for inserted DNA, and those labeled +/+ are WT. M indicates size markers, and BAC indicates a genomic BAC clone that includes the *lnx2a* locus.

Our interpretation of these results is illustrated in Figure 6. Donor DNA was inserted into the genome of some of the injected fish at an unknown locus, not in the target region within the *lnx2a* gene. This presumably occurred by transgene insertion in a manner that was generally employed before more efficient methods were introduced, such as meganuclease enhanced or Tol2 mediated transgenesis [25,26]. When DNA from these transgenic fish was used as template in PCR reactions, using primer pairs that bridge genomic sequence outside the homology arms with donor sequence, recombination occurred *in vitro*, as described by Meyerhans and colleagues more than 25 years ago [1]. PCR reactions start at their appropriate location in different regions of the genome. When elongation intermediates containing complementary sequences within the homology arms fall off the template they can hybridize, restart elongation, and then act as template for standard PCR amplification. While such *in vitro* recombination by hybridization of incomplete products may be a rare event, the extraordinary efficiency of PCR means that rare events can lead the robust outputs. The *in vitro* product will then have precisely the same sequence that is predicted for the product of *in vivo* homologous recombination.

**Fig. 6.**
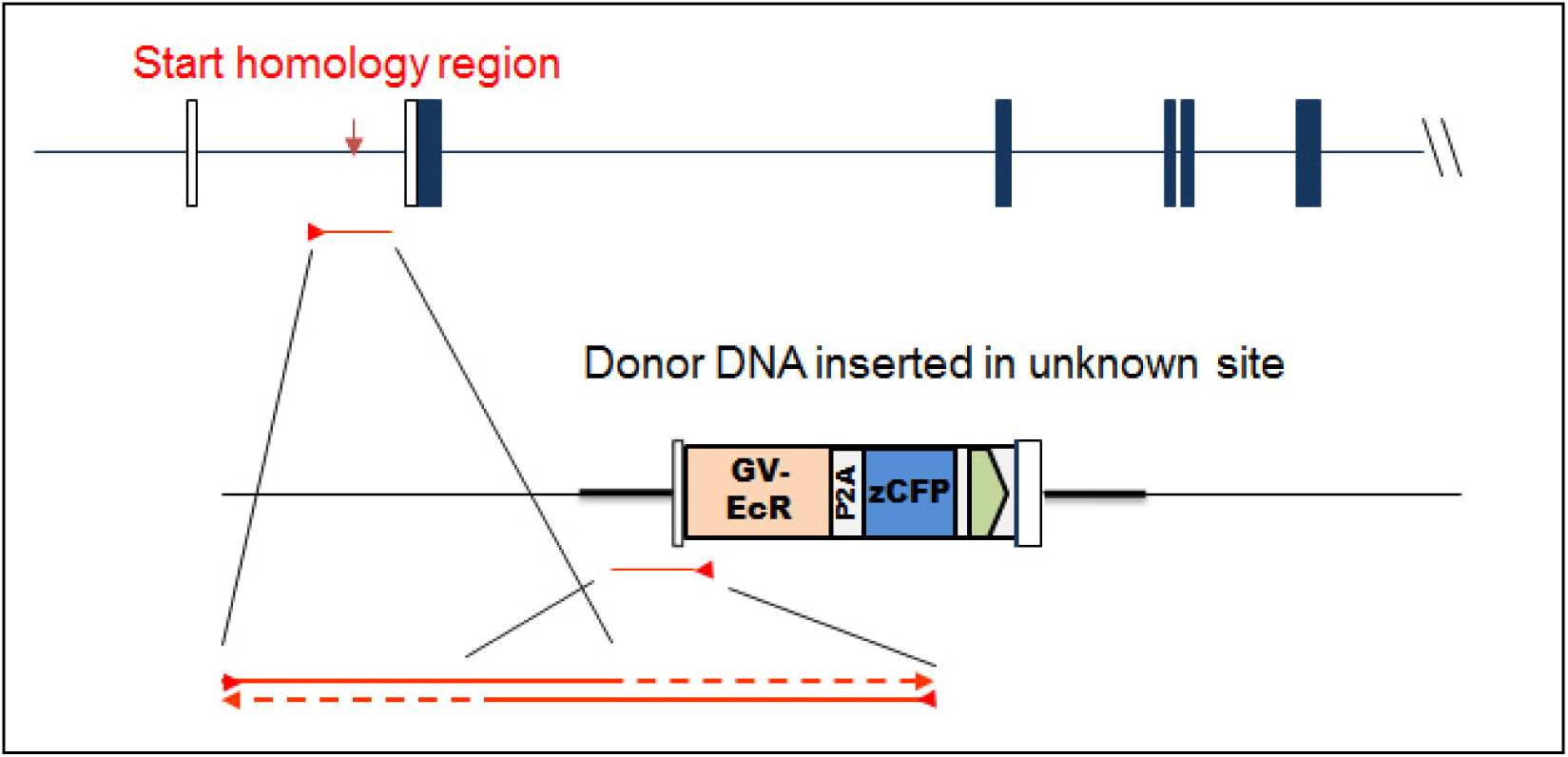
Model for in vitro recombination during PCR procedure. Based on the Southern blots (Fig. 5) and earlier reports of *in vitro* recombination (Meyerhans 1990), we propose the following interpretation. In certain injected fish the donor DNA was incorporated into an unknown position of the genome. PCR reactions started in the *lnx2a* genomic locus and in the transgene, as indicated. When molecules stopped elongation and fell off the template anywhere in the homology region, overlapping ends of complementary products could anneal and restart elongation, generating a recombinant molecule *in vitro.* Such molecules could then be amplified by the primer pair used in the reaction.

## Conclusions

With the popularity of genome editing by homologous recombination in zebrafish increasing [11-18] we wish to point out a potential artifact in PCR-based testing for recombinant products in such experiments. We also would like to draw attention to findings reported long ago that established the potential for *in vitro* recombination during PCR reactions under conditions where products with overlapping complementary sequences may arise [1]. We would like to stress that we do not question the validity of any one individual report of homologous recombination in zebrafish, we simply want to relate our experience of a compelling-looking artifact. We believe that methods in addition to PCR should be used in verifying homologous recombination, as indeed have been used in most of the literature to date. We also would like to stress the value of Southern blots for verifying certain kinds of genome editing – the lower sensitivity of this method as compared to PCR is in fact an asset in this context.

## Materials and Methods

Please refer to our earlier publication [19] for most of the materials and methods used. See Supplementary Information Figures S1 and S2 for sequences of Donor DNA and homology arms.

### Genomic PCR

Genomic DNA was isolated from embryos by the HotSHOT method [27,28]. PCR was performed using AccuPower PCR Premix (Bioneer Inc.).

Primers used in Fig. 3B and C: 5’ PCR (F-CACCATCTTAAAACGTTTACTGTGT, R-ACTTGGCGCACTTCGGTTTT); 3’ PCR (F-CGTATTACGTAGATCCAGACATGAT,

R-TGAACATGTTTAAAGCACAGACCT); CFP PCR (F-ACTGGAGTCGTGCCTATCCT, R-CTGCTTCATGTGGTCAGGGT).

## Plasmid construction

Zebrafish Lig4-DN (N-terminal 655aa deletion) and Ku70-DN (N-terminal 57aa deletion) used in injection experiments, was amplified from zebrafish cDNA using RT-PCR, following the approach reported for human genes [29,30]. PCR products were cloned into the pCS2+MT expression vector between XhoI and XbaI sites. Primers and sequences of these products are given in Supplementary Figures S3 and S4.

## Southern blot analysis

Southern blot was performed using 10μg of genomic DNA of individual F1 fish. Total genomic DNA was purified by proteinase K digestion in buffer containing 10 mM Tris (PH 8.0), 100 mM EDTA (PH 8.0), 0.5% SDS, and 400 µg/mL proteinase K, followed by phenol extraction. The DNA was digested overnight with Hind III, precipitated with ethanol, separated on a 0.8% agarose gel, and transferred onto a positively charged nylon membrane. The DNA was UV crosslinked to the membrane, and hybridized with DIG-labeled probes (Roche, DIG High Prime DNA Labeling and Detection Starter Kit II).

## Acknowledgement

We thank Harry Burgess for encouragement and valuable advice on the manuscript, and for providing reagents. This work was supported by the Intramural Research Program of the National Institute of Child Health and Human Development, National Institutes of Health.

